# Context-Aware Protein Representations Using Protein Language Models and Optimal Transport

**DOI:** 10.64898/2026.01.24.701517

**Authors:** Sahil Patel, Navid NaderiAlizadeh

**Author notes:** { }.

## Abstract

Proteins have different functions in different contexts. As a result, representations that take into account a protein’s biological context would allow for a more accurate assessment of its functions and properties. Protein language models (PLMs) generate amino-acid-level (residue-level) embeddings of proteins and are a powerful approach for creating universal protein representations. However, PLMs on their own do not consider context and cannot generate context-specific protein representations. We introduce COPTER, a method that uses optimal transport to pool together a protein’s PLM-generated residue-level embeddings using a separate context embedding to create context-aware protein representations. We conceptualize the residue-level embeddings as samples from a probabilistic distribution, and use sliced Wasserstein distances to map these samples against a context-specific reference set, yielding a contextualized protein-level embedding. We evaluate COPTER’s performance on three downstream prediction tasks: therapeutic drug target prediction, genetic perturbation response prediction, and TCR-epitope binding prediction. Compared to state-of-the-art baselines, COPTER achieves substantially improved, near-perfect performance in predicting therapeutic targets across cell contexts. It also results in improved performance in predicting responses to genetic perturbations and binding between TCRs and epitopes. The implementation code is available at https://github.com/SahilP113/COPTER.

## 1. Introduction

Proteins are the primary functional macromolecules of the cell, executing a wide range of biological processes including catalysis, molecular recognition, signal transduction, and immune response [1, 14, 43]. A protein’s biological activity is governed by its sequence, structure, and interactions, which together determine its function across different cellular and environmental contexts. As a result, learning informative and expressive protein representations is a central problem in computational biology. High-quality protein representations enable a broad range of downstream prediction tasks including protein function annotation, structure and interaction prediction, therapeutic target discovery, and modeling cellular responses to perturbations [29, 60].

One of the most powerful emerging classes of protein representation learning methods is protein language models (PLMs), large-scale pre-trained foundation models that generate protein embeddings from amino acid sequences. Using self-supervised learning, PLMs are trained on massive collections of protein sequence data and learn representations that capture evolutionary constraints and sequence-level regularities. PLMs have demonstrated remarkable predictive ability across a wide range of sequence-based prediction tasks [25, 31, 40, 52].

Given a protein sequence of length *n*, a PLM produces an embedding in ℝ^*n*×*d*^ where *d* denotes the dimensionality of the PLM latent embedding space. Each amino acid embedding, or token-level embedding, not only captures information about the corresponding residue but also contextual information about the entire protein sequence. The most common method to aggregate these token-level embeddings into a single representation for the entire protein is average pooling [2, 52, 53, 58]. In average pooling, the mean of all token-level embeddings is obtained to produce a fixed-length pooled representation in ℝ^*d*^. A limitation of average pooling is that it treats each amino acid embedding equally. This is often inaccurate, as a subset of amino acids is often primarily responsible for a protein’s structure, interactions, and function.

To address this limitation, recent approaches have been proposed to construct more informative global protein representations from residue-level embeddings. In [37], an optimal transport-based method was introduced to pool amino acid-level PLM embeddings. This approach learns directions, or slices, in ℝ^*d*^ onto which both the input embeddings and a set of learnable reference embeddings are projected. Based on these projections, optimal transport couplings between the input and reference empirical distributions are used to derive a fixed-length protein-level representation.

Nevertheless, proteins operate within specific biological contexts, such as the cell type or physiological condition in which they are expressed. A protein’s function, interactions, and regulatory roles can vary substantially across contexts, even when its amino acid sequence remains unchanged [19, 41]. As a result, incorporating contextual information into protein representations has the potential to enable more accurate prediction of protein properties and interactions [5, 16, 42]. However, the vast majority of existing protein representation learning approaches, including PLMs, produce context-agnostic embeddings that assign a single fixed representation to a protein regardless of the biological context in which it is observed.

The usefulness of contextualized protein representations has been demonstrated across a wide range of downstream tasks, including drug-target identification, prediction of responses to genetic and chemical perturbations, protein-peptide interaction prediction, and cross-species protein function prediction [29, 34, 59, 60]. Together, these applications highlight the flexibility of the notion of context, which need not be limited to cellular context but can also encompass interacting proteins, gene expression profiles, and other relevant biological signals.

In this work, we introduce a framework that directly incorporates biological context into the aggregation process of frozen residue-level protein language model embeddings to produce context-aware protein-level representations. By integrating biological context into the pooling process, our approach enables the construction of protein representations that adapt to the context in which a protein is observed and more accurately capture context-dependent properties and interactions. Our method, which we refer to as **COntextualized ProTein Embeddings via optimal tRansport (COPTER)**, combines optimal transport–based pooling with context-specific embeddings to derive context-aware protein representations. By jointly leveraging the expressive power of PLMs and the flexibility of optimal transport, our framework provides a principled and robust approach for modeling proteins across diverse biological contexts. To the best of our knowledge, COPTER is the first approach to integrate contextual information directly into the aggregation process of residue-level PLM embeddings.

We evaluate COPTER across three downstream tasks: therapeutic drug target prediction, gene expression response prediction to genetic perturbations, and T-cell receptor (TCR)-epitope binding prediction. Incorporating contextual information into protein representations leads to substantial performance gains in therapeutic target prediction and more modest, yet consistent, improvements in genetic perturbation response and TCR-epitope binding prediction. These results demonstrate the value of context-aware protein representations for modeling protein function across diverse biological settings. More broadly, COPTER provides a flexible framework that can be extended to incorporate additional sources of biological context, opening avenues for future applications in protein interaction modeling, functional genomics, and precision medicine.

## 2. Related Work

### 2.1. Protein Language Models

Protein language models (PLMs) are inspired by large language models (LLMs) and capture the language of life written into protein sequences. These models are trained primarily on protein sequence data using the masked language modeling (MLM) objective, predicting masked amino acids based on their surrounding amino acids. Many prominent PLMs exist, including ESM-2 [30, 48], ProtTrans [10], ProtFluentE1 [20], and ProGen3 [3].

The residue-level embeddings produced by PLMs have been used in a wide variety of downstream tasks, including peptide generation [4], protein engineering [13], post-translational modification prediction [44], predicting protein-protein interactions [57], prediction of secondary structural properties [50], antibody design [63], and allosteric binding site prediction [47].

### 2.2. Pooling Protein Language Model Representations

PLMs produce embeddings at the amino acid, or residue, level for a given input sequence. Because protein sequences vary in length, the number of residue-level embeddings differs across proteins, necessitating aggregation methods that map variable-length sets of embeddings to fixed-length protein-level representations.

The most common aggregation strategy is average pooling, which satisfies both permutation and size invariance. Other simple pooling operations, such as max pooling and softmax pooling, have also been explored [51]. More recently, several learned pooling methods have been proposed to better capture informative residues, including locality-aware [18], attention-based [55], and PageRank-based pooling strategies [56].

Among these approaches, optimal transport (OT)-based aggregation methods provide a principled mechanism for pooling residue-level embeddings by modeling them as distributions and aligning them to learnable reference sets [36, 37]. In this work, we build on such OT-based aggregation techniques and extend them by incorporating biological context directly into the pooling process.

### 2.3. Contextual Protein Representation Learning

Incorporating information about a protein’s context into its representation could enable more accurate predictions of its functions and properties. The increasing prevalence of single-cell gene expression databases has made it much easier to assess how gene expression, and as a result, protein expression, differs based on context [5, 27, 67]. This has led to the development of machine learning models that train on context-specific data and learn contextualized protein representations. For example, a geometric deep learning model called PINNACLE was developed to create contextualized protein representations by training on a network of protein interactions and a metagraph of cellular interactions, learning a latent representation space based on the position and cell context of proteins [29]. Other deep learning methods have also been developed to learn contextualized protein embeddings for downstream prediction tasks such as ProteomeLM [34] and AggrescanAI [38].

The increasing popularity of contextual learning led to the creation of Therapeutic Data Commons 2 (TDC-2) [60]. TDC-2 provides a platform that integrates single-cell analysis with multimodal machine learning through four contextual AI tasks. We use the datasets and benchmarks provided by TDC-2 to evaluate the performance of our method on three contextual tasks: therapeutic-target nomination, TCR-Epitope binding prediction, and genetic perturbation response prediction.

## 3. Methods

### 3.1. Problem Formulation

Consider a protein sequence **s** = (*s*_1_, *s*_2_, …, *s*_*n*_) ∈ ℙ^*n*^ comprising *n* residues, where ℙ denotes the alphabet of 20 canonical amino acids. In addition, consider a context vector in **c** ∈ ℝ^*k*^ which represents the biological context in which the protein is found. The overall goal of context-aware protein representation learning is to find a *context-aware embedding* function *e*_*θ*_ : ℙ^*n*^ × ℝ^*k*^ → ℝ^*L*^, potentially parameterized by a set of parameters denoted by *θ*, that maps the protein sequence **s** to a context-aware *L*-dimensional latent embedding *e*_*θ*_(**s, c**). Note that the dimensionality of the embedding space, *L*, is independent of the length of the protein sequence, so a protein sequence of any length needs to be mapped to a fixed-dimensional embedding in ℝ^*L*^.

As an integral component of the context-aware embedding function *e*_*θ*_, we leverage a protein language model (PLM), which can be denoted by a function *f*_PLM_ : ℙ^*n*^ → ℝ^*n*×*d*^, mapping every amino acid to an embedding in ℝ^*d*^. A pooling function can then be applied to the PLM output to aggregate the amino-acid level embeddings into a single representation for the entire protein in ℝ^*L*^.

Our goal in this paper is to incorporate the context information **c** into the pooling operation and design a *context-aware pooling* function *p*_*θ*_ : ℝ^*n*×*d*^ × ℝ^*k*^ → ℝ^*L*^ so that the resulting protein-level representation in ℝ^*L*^ takes the context into account. The end-to-end context-aware embedding function *e*_*θ*_ is then given by the composition of the PLM and context-aware pooling function, i.e., *e*_*θ*_(**s**) := *p*_*θ*_(*f*_PLM_(**s**)). Note that the PLM parameters are kept frozen, implying that we only parameterize and train the context-aware pooling function *p*_*θ*_.

### 3.2 Context-Aware Pooling via Optimal Transport (COPTER)

We propose a context-aware pooling mechanism that integrates biological context directly into the aggregation of residue-level PLM embeddings. Our approach extends prior optimal transport-based pooling methods, specifically sliced Wasserstein embedding (SWE) pooling [9, 26, 37], by conditioning the pooling operation on the protein’s context vector.

Given a protein sequence **s** ∈ ℙ^*n*^, we first obtain residue-level embeddings *f*_PLM_(**s**) ∈ ℝ^*n*×*d*^ using a pretrained PLM with frozen parameters. In SWE pooling, these residue-level embeddings are treated as samples from an underlying empirical distribution in ℝ^*d*^. SWE aggregates embeddings by projecting both the input embeddings and a set of learnable reference points onto multiple one-dimensional subspaces (“slices”) and computing optimal transport couplings in each slice. In the original formulation, the reference points are learned globally and are shared across all inputs. Our key idea is to make these reference points context-dependent, thereby allowing the pooling operation itself to adapt to the biological context.

Formally, let *L* and *m* denote the number of slicing directions and reference points, respectively. We design a context-aware pooling function *p*_*θ*_ by conditioning the sliced SWE reference embeddings on the context vector **c** ∈ ℝ^*k*^. Specifically, we use a linear hypernetwork [17], represented by *h*_*ϕ*_ : ℝ^*k*^ → ℝ^*m*×*L*^, to map the context vector **c** to sliced reference embeddings *h*_*ϕ*_(**c**) ∈ ℝ^*m*×*L*^. This construction ensures that contextual information is injected directly into the reference distribution that guides the optimal transport alignment.

In parallel, we project the residue-level embeddings *f*_PLM_(**s**) ∈ ℝ^*n*×*d*^ through *L* trainable linear slicing operators, producing *d* empirical one-dimensional distributions. For each slice, we compute the Monge coupling between the projected residue embeddings and the corresponding context-aware sliced reference points using sorting and interpolation operators as in [37]. These couplings yield *L* vectors in ℝ^*m*^, which are concatenated to form a matrix in ℝ^*L*×*m*^. A final learnable projection aggregates this matrix into a fixed-length protein embedding in ℝ^*L*^, yielding the final context-aware protein representation. Figure 1 provides an overview of our approach.

**Figure 1.**
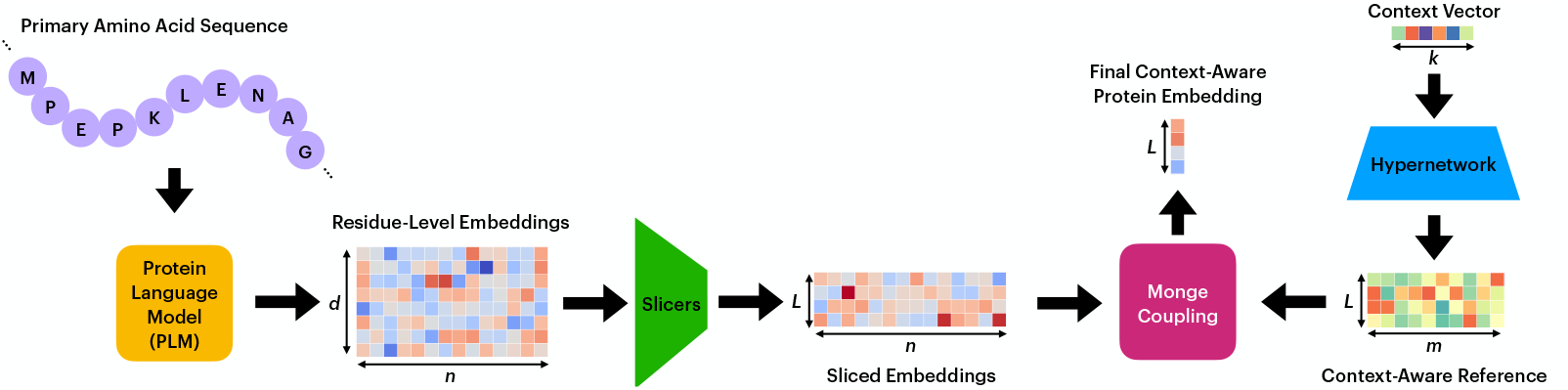
: Overview of COPTER. A protein’s primary amino acid sequence is first encoded by a pretrained protein language model (PLM) to produce residue-level embeddings. These embeddings are projected onto multiple one-dimensional subspaces using learnable slicing directions. In parallel, a biological context vector is passed through a hypernetwork to generate a context-aware reference distribution. COPTER computes sliced optimal transport (Monge) couplings between the projected residue embeddings and the context-conditioned reference, and aggregates these couplings to produce a final context-aware protein embedding.

The resulting context-aware protein embeddings are fed into a task-specific multi-layer perceptron (MLP) for downstream prediction. The context-aware COPTER pooling module and the MLP are trained jointly, while the PLM remains frozen. The form of the context vector **c** depends on the downstream task. For therapeutic target prediction, context embeddings are derived from protein interaction graphs using a graph neural network. For predicting genetic perturbation responses, the context vector corresponds to the cell’s pre-perturbation gene expression profile. For TCR-epitope binding prediction, the context is represented by mean-pooled embeddings of the epitope sequence obtained from a PLM. Despite these differences, COPTER provides a unified mechanism for incorporating heterogeneous biological context into protein representations.

### 3.3. Protein Language Models and Embedding Extraction

We evaluate COPTER using multiple pretrained PLMs to assess robustness across architectures and model scales. Specifically, we use ESM-2 with 8 million parameters (ESM2-8M) and 650 million parameters (ESM2-650M), as well as ProGen2-small and ProGen2-large. For ESM2-8M and ESM2-650M, the embedding dimension *d* is 320 and 1280, respectively, while ProGen2-small and ProGen2-large, respectively, produce embeddings of dimension 1024 and 2560. All PLMs are used with frozen parameters, and only the context-aware pooling module and downstream prediction models are trained. This setup allows us to isolate the effect of contextual pooling and evaluate COPTER independently of PLM fine-tuning.

### 3.4. Therapeutic Target Prediction

We reproduced the 156 context-specific protein interaction networks and the metagraph of cell and tissue embeddings used by PINNACLE [29]. Using the same parameters, we fed the protein interaction networks and the metagraph into the PINNACLE geometric deep learning model. PINNACLE produced cell embeddings for each of the 156 cell contexts, tissue embeddings, and protein representations for each cell context. We used the 64-dimensional cell embeddings produced by PINNACLE as the cell context representations in this task. The performance of the contextualized protein embeddings produced by PINNACLE served as a baseline for COPTER.

We also used the same proteins employed to construct the context-aware protein interaction networks in this task. We fed their amino acid sequences into a pretrained PLM, specifically ESM2-8M, ESM2-650M, ProGen2-small, or ProGen2-large. For each protein in each cell context, we aggregated the PLM’s token-level embeddings with the cell context representation produced by PINNACLE using COPTER. As a result, for each protein, we generated a separate representation for every cell in which it was found. Each representation was a 64-dimensional vector, corresponding to the cell context’s dimensionality.

We then fed the contextualized protein representations into an MLP. We jointly trained the COPTER pooling model and the MLP classifier to predict therapeutic targets for rheumatoid arthritis (RA) and inflammatory bowel disease (IBD). The task involved binary classification of whether a given protein in a specific cell context is a therapeutic target.

### 3.5. Genetic Perturbation Response Prediction

The purpose of the genetic perturbation response prediction task is to predict the effects of perturbing one or more genes in a cell. In particular, it predicts how perturbing one or more genes in a cell affects the expressions of all the other genes in the cell. Previous machine learning models have been developed for predicting responses to genetic perturbations [8, 32, 49]. We used COPTER for the genetic perturbation prediction task by considering the amino acid sequence of the protein encoded by the perturbed gene and using the pre-perturbation gene expression as the context.

We performed this task using three different genetic perturbation datasets: Replogle RPE1 [46], Replogle 562 [46], and Norman K562 [39]. The Replogle RPE1 and Replogle 562 datasets were pre-processed to include 5000 genes, while the Norman K562 dataset was preprocessed to include 5045 genes, following the procedure used by the state-of-the-art GEARS method [49]. The Replogle RPE1 and Replogle 562 datasets included only single genetic perturbations, meaning that only one gene was perturbed in each cell. The Norman K562 dataset included single and double perturbations, allowing up to 2 genes to be perturbed per cell.

For each perturbed gene, the amino acid sequence of the primary isoform encoded by the gene was obtained and fed into a PLM, resulting in a sequence of amino acid embeddings. The context embedding was the vector of gene expressions in the cell prior to perturbation. We used COPTER to pool amino acid embeddings for the main isoform using the gene expression context. For the double perturbations, COPTER was independently applied to each perturbation, yielding a pooled vector for both. These two vectors were then summed to obtain an embedding that represented the combined effect of both perturbations.

We set *L* = 320 for all genetic perturbation prediction tasks. As a result, COPTER produced a 320-dimensional vector. The MLP projected this 320-dimensional embedding to a *k*-dimensional embedding using two hidden layers. This embedding represented the predicted *change* in gene expression due to the perturbation. The mean squared error loss between the predicted change in gene expression vector and the true change in gene expression vector was calculated for the top 20 most differentially-expressed genes. The true perturbation effects were determined by finding the difference in gene expression between a specific perturbed cell and a randomly sampled control cell.

### 3.6. TCR-Epitope Binding Prediction

The TCR-epitope binding task aims to predict whether an epitope will bind to a T-cell receptor (TCR) [60, 62]. This task also involves binary classification of whether the TCR and epitope will bind. Many deep learning approaches have been developed to predict TCR-epitope binding [6, 23, 24, 54, 62]. Our approach considers the amino acid sequences of both the TCR and the epitope, using the TCR as the protein and the epitope as the context.

We used three different methods to generate negative samples: random shuffling of epitope and TCR sequences (RN), experimental negatives (NA), and pairing external TCR sequences with epitope sequences (ET) [60]. For the RN and NA tasks, we obtained data from the TC-hard dataset, and for the ET task, we obtained data from the PanPep dataset [12]. For the RN and NA heuristics, the CDR3*α* and CDR3*β* chains were included in the TCR sequence, whereas the TCR sequence for the ET method included only the CDR3*β* chain. We used the TCR sequence as the query sequence (comprising either both the CDR3*α* and CDR3*β* chains or only the CDR3*β* chain), which we fed into a PLM.

For all three tasks, we used the epitope sequence as the context. In particular, we also fed the epitope sequence into the same PLM as the TCR. Given the short length of the epitope sequence [37], we used average pooling for the epitope token-level embeddings, resulting in a single vector in ℝ^*d*^ for the epitope embedding. This pooled *d*-dimensional vector was used as the context.

We used COPTER to aggregate token-level embeddings of the TCR sequence, conditioned on the epitope embedding as the context, to create a contextualized TCR representation. Since the size of the context (epitope embedding) was *d*, which depended on the PLM, the resulting contextualized representation was a *d*-dimensional vector. We set *L* = *d* to ensure equivalence with the context size. The contextualized TCR representation was then passed into the final MLP classifier, which predicted the probability of TCR-epitope binding.

## 4. Results

### 4.1. Therapeutic Target Prediction

Table 1 compares the performance of COPTER on the RA and IBD tasks with that of PINNACLE and a graph attention network (GAT) [61] baseline across five different random seeds. COPTER substantially outperforms both GAT and PINNACLE, achieving near-perfect performance with respect to context-free AP@5 and AUROC. This holds true across all four PLMs and for both the RA and IBD tasks. Additionally, our results show that COPTER exhibits near-perfect performance when evaluated using context-specific metrics, including AP@5 for the top 20 cell types and AUROC for top-1, top-10, and top-20 cell types. This demonstrates that the protein representations learned by COPTER are robust both within specific cell contexts and when considered independently of context.

**Table 1:** Performance of COPTER compared to PINNACLE and GAT on the RA and IBD therapeutic target prediction tasks, reported in terms of mean and standard deviation across five different random seeds. Best results are highlighted in **bold**. AP@5 CF: Context-free AP@5 integrated across all cell types. AUROC CF: Context-free AUROC integrated across all cell types.

| Model |  | Task | AP@5 CF (↑) | AUROC CF (↑) |
| --- | --- | --- | --- | --- |
| PINNACLE |  | RA | 0.226 ± 0.023 | 0.510 ± 0.005 |
| GAT |  |  | 0.220 ± 0.013 | 0.580 ± 0.010 |
| COPTER | ESM2-8M |  | **1.000 ± 0.000** | 0.997 ± 0.001 |
|  | ESM2-650M |  | **1.000 ± 0.000** | 0.996 ± 0.001 |
|  | ProGen2-Small |  | **1.000 ± 0.000** | **0.998 ± 0.000** |
|  | ProGen2-Large |  | **1.000 ± 0.000** | 0.994 ± 0.001 |
| PINNACLE |  | IBD | 0.198 ± 0.013 | 0.500 ± 0.010 |
| GAT |  |  | 0.200 ± 0.023 | 0.640 ± 0.017 |
| COPTER | ESM2-8M |  | **1.000 ± 0.000** | **0.999 ± 0.000** |
|  | ESM2-650M |  | **1.000 ± 0.000** | **0.999 ± 0.001** |
|  | ProGen2-Small |  | **1.000 ± 0.000** | **0.999 ± 0.001** |
|  | ProGen2-Large |  | **1.000 ± 0.000** | 0.997 ± 0.001 |

Furthermore, Figure 2 provides a t-SNE [33] visualization of the protein embeddings created by COPTER with all four PLMs on the RA task, where colors represent cell context. These protein embeddings are all generated in a zero-shot manner (i.e., using randomly initialized COPTER parameters and without downstream classification training), demonstrating COPTER’s inherent ability to separate protein embeddings based on the cellular context.

**Figure 2.**
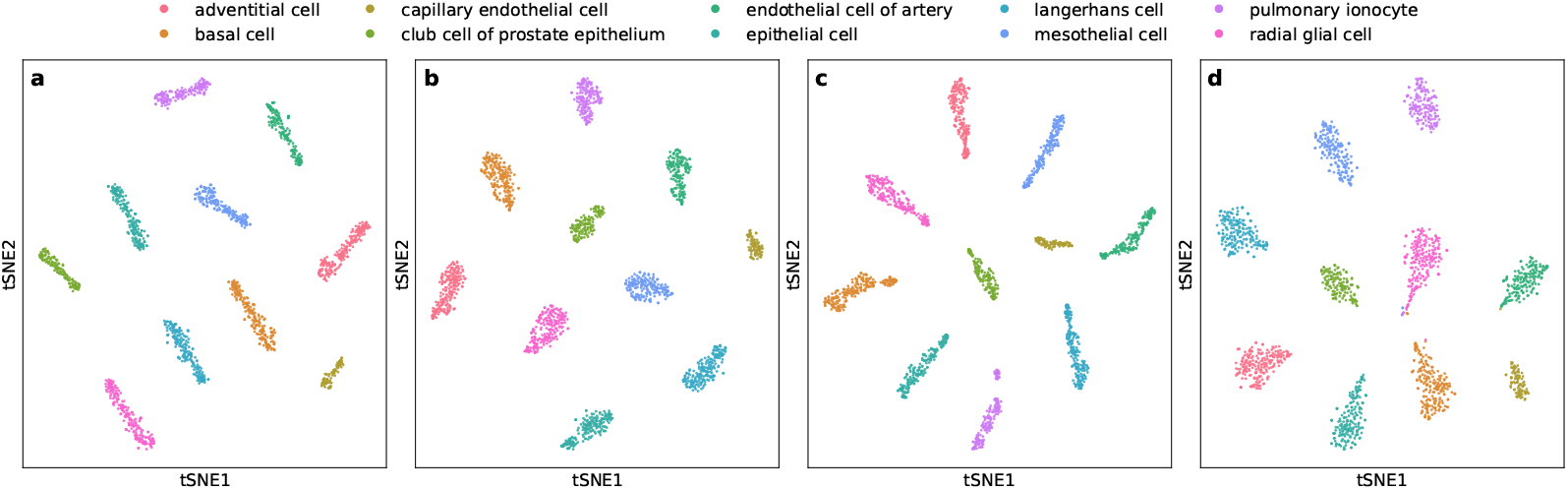
: t-SNE visualizations of protein embeddings generated by COPTER for the RA task across 10 randomly-selected cell types, colored accordingly, using SWE pooling with (a) ESM2-8M, (b) ESM2-650M, (c) ProGen2-small, and (d) ProGen2-large.

### 4.2. Genetic Perturbation

Table 2 compares the performance of COPTER across five different random seeds to that of genetic perturbation baseline models, including GEARS [49], a deep-learning based model called CPA [32], and a baseline which assumes gene expression remains unchanged after perturbation (no-perturb).

**Table 2:**
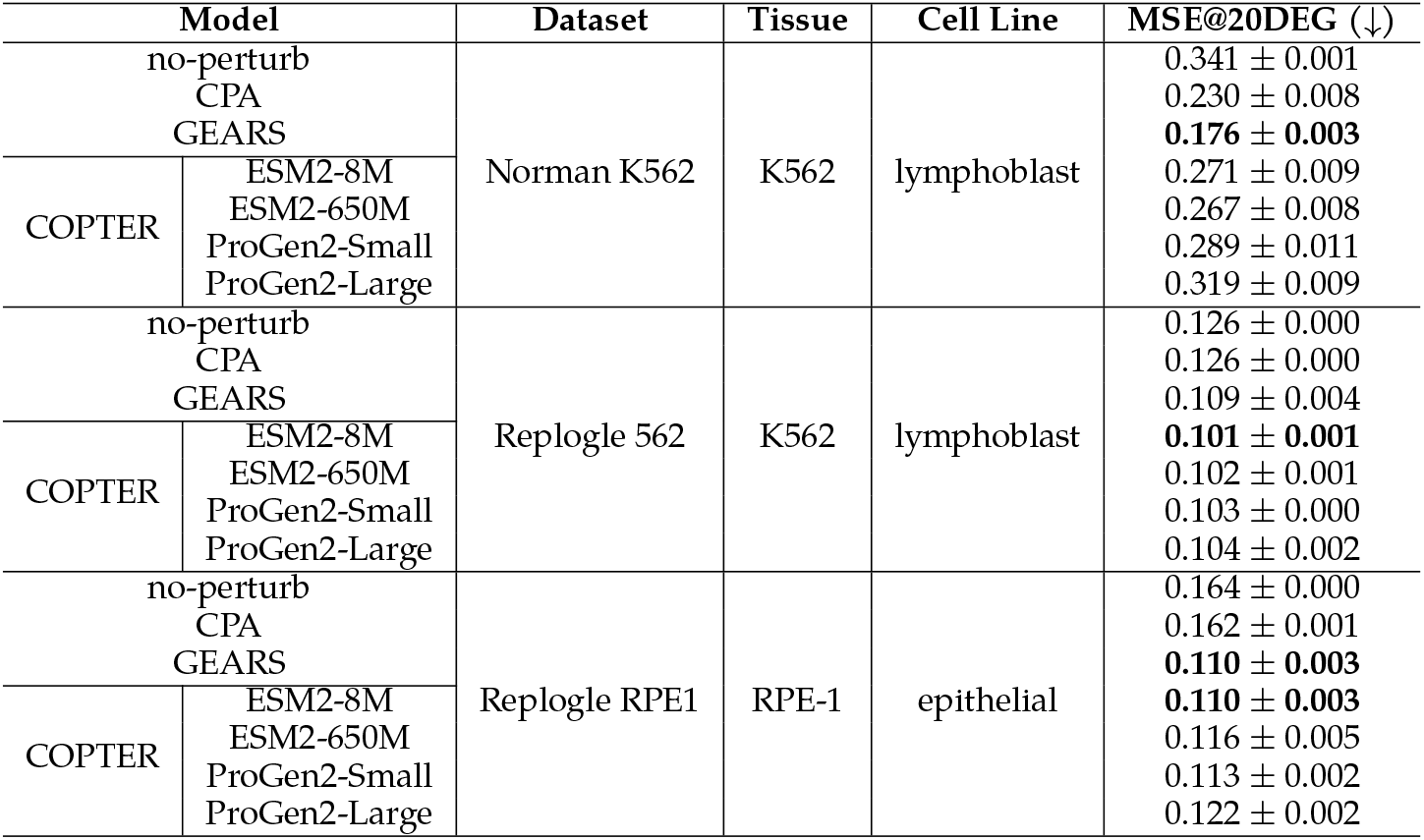
Mean and standard deviation of COPTER’s performance across five different random seeds compared to baselines on three genetic perturbation datasets. Best results are highlighted in **bold**. MSE@20DEG: Mean squared error of the top 20 most differentially expressed genes.

When evaluated on the Norman dataset (which includes both single- and double-gene perturbations), COPTER has a higher MSE than GEARS across all four PLMs. The best-performing COPTER method on this dataset, which used the ESM2-650M PLM, produced a mean MSE higher than those of GEARS and CPA. We also assessed COPTER’s performance separately on single and double perturbations from this dataset. COPTER exhibited poorer performance than GEARS across both single- and double-gene perturbations.

However, all four COPTER methods showed improved performance in predicting responses to genetic perturbations on the Replogle 562 dataset. Furthermore, on the Replogle RPE1 dataset, COPTER generally achieved very similar performance to GEARS. More specifically, using ESM2-8M yielded the same MSE as GEARS on the Replogle RPE1 dataset, while the other three PLMs performed marginally worse.

These results demonstrate that COPTER’s success in predicting responses to genetic perturbations depends on the dataset. COPTER appears to perform better when trained and evaluated on single-gene perturbations.

### 4.3. TCR-Epitope Binding Prediction

Table 3 compares the performance of COPTER with that of state-of-the-art methods for TCR-Epitope binding prediction including AVIB-TCR [15], MIX-TPI [64], Net-TCR2 [21], PanPep [12], TEINet [22], and TITAN [62]. across the RN, NA, and ET tasks.

**Table 3:**
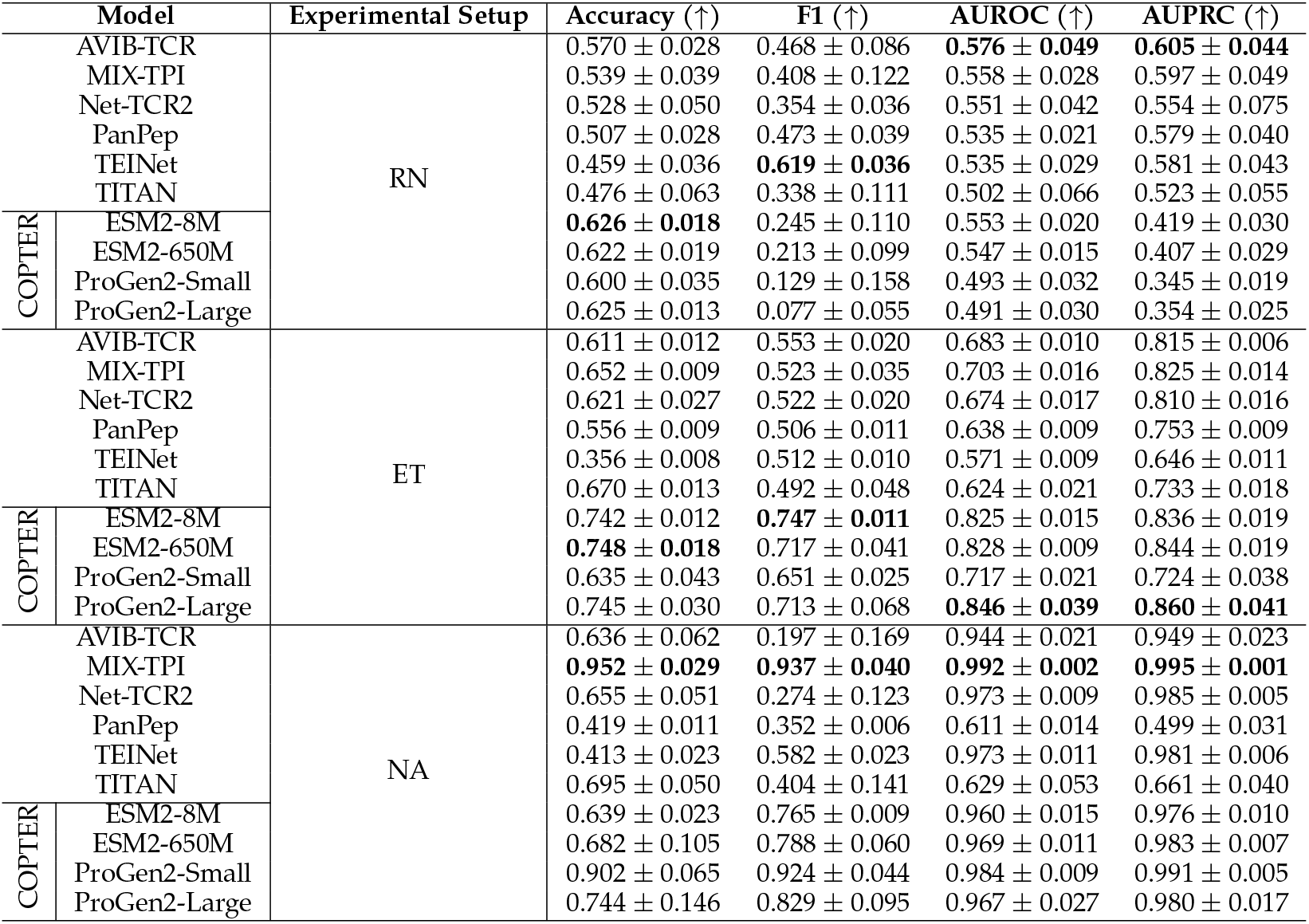
Five-fold cross-validation performance of COPTER on the RN, NA, and ET tasks compared to baseline models. Best results are highlighted in **bold**.

For the RN task, COPTER performed best with the two ESM2 PLMs. COPTER with ESM2-8M out-performed all methods in accuracy. However, its F1 score and its AUPRC underperformed those of all baseline methods. The AUROC of COPTER fell between the baseline methods, outperforming four of them but performing worse than the other two. Even with the ESM2 PLMs, while COPTER shows greater accuracy than the baseline methods, it does not have an overall advantage over them due to its lower F1 and AUPRC scores.

The best performing model for the NA task was MIX-TPI. However, the four COPTER models generally outperformed all other baseline methods. COPTER exhibited the best performance with the ProGen2-small PLM, achieving higher scores across all metrics relative to the other PLMs.

For the ET task, COPTER with ESM2-8M outperformed all six traditional methods across all four metrics. The other COPTER models also outperformed all baseline methods. Therefore, COPTER exhibited a very strong performance on the ET task.

## 5. Discussion

We introduced COPTER, a new method for contextualizing protein language model representations via optimal transport. COPTER uses a PLM to generate residue-level embeddings and optimal transport to pool them into a contextualized protein representation, conditioning on the context embedding as side information.

The contextualized protein representations generated by COPTER substantially outperformed PINNACLE-generated protein embeddings in predicting therapeutic drug targets, achieving near-perfect performance. This improvement in performance held true for both the RA and IBD tasks. The particular success of the contextualized representations on the therapeutic target prediction task is most likely because using a PLM to create amino-acid embeddings effectively represents residue-level information about the protein. Maintaining residue-level information is especially important when predicting whether a protein binds to a drug, since the interactions of amino acids with the drug will determine binding. The zero-shot protein embeddings produced by COPTER were very well separated by cell context for both the RA and IBD tasks. COPTER’s ability to separate protein embeddings based on their context in a zero-shot manner is supported by previous work with optimal transport pooling, which found that assigning random parameters to the SWE model without training still resulted in strong performance [37].

COPTER produced mixed results for predicting genetic perturbation responses, performing slightly better than GEARS on the Replogle 562 dataset, similar on the Replogle RPE1 dataset, and noticeably worse on the Norman dataset. A limitation of the genetic perturbation task is that only the primary isoform of the perturbed gene was considered. This main isoform was used as the protein representation of the gene. Due to alternative splicing, a gene can code for multiple proteins, resulting in multiple isoforms that represent the gene. Therefore, only considering the main isoform may not be sufficient when assessing the effects of a gene perturbation.

For predicting TCR-epitope binding, COPTER generally demonstrated improved performance, though the improvements it achieved were smaller than those observed for therapeutic target prediction. TCR-epitope binding prediction is an inherently challenging task [60]. Negative samples (non-binders) are underrepresented in TCR-epitope binding datasets and heuristics must be used to generate more negative samples. These heuristics, such as RN, NA, and ET, often lead to unsatisfactory model performance. The inherent difficulty of successfully predicting TCR-epitope binding with these heuristics may limit the amount of improvement that can be made on these tasks.

For future work, more parameters could be varied in the SWE pooling method. In all experiments, the numbers of slicers and reference points were kept constant. Varying the number of slicers and reference points could potentially improve protein representations. More recent PLM families, such as ESM-C [11], ProGen3 [3], and E1 [20], could be evaluated to assess whether newer training paradigms yield improved contextualized downstream performance. Additional tasks could also be carried out to determine the effectiveness of COPTER for representing proteins. For example, these representations could be used to predict responses to chemical perturbations [45, 60, 65], predict enzyme-substrate binding [7, 28], or study ligand-GPCR interactions [66] Furthermore, for the therapeutic target prediction tasks, the GNN used to create the cell-context embeddings was kept frozen. However, the GNN parameters could also be finetuned during SWE pooling training to produce improved cell-context embeddings. This fine-tuning can also be extended to PLM parameters rather than keeping them frozen, as prior work has shown the benefits of fine-tuning PLMs across a variety of protein representation learning tasks [35, 50, 53].

Overall, COPTER provides a principled framework for learning protein representations that integrate residue-level information with biological context. By incorporating contextual signals directly into the aggregation of protein language model embeddings, COPTER produces more expressive and adaptive protein representations, enabling improved modeling of protein function and interactions across diverse biological settings.

